# Ubiquitylation of nucleic acids by DELTEX ubiquitin E3 ligase DTX3L

**DOI:** 10.1101/2024.04.19.590267

**Authors:** Kang Zhu, Chatrin Chatrin, Marcin J Suskiewicz, Vincent Aucagne, Dragana Ahel, Ivan Ahel

## Abstract

Recent discoveries expanding the spectrum of ubiquitylation substrates to include non-proteinaceous molecules have broadened our understanding of this modification beyond conventional protein targets. However, the existence of additional types of substrates remains elusive. Here, we present evidence that nucleic acids can also be directly ubiquitylated. DTX3L, a member of the DELTEX family E3 ubiquitin ligases, ubiquitylates DNA and RNA *in vitro* and that this activity is not shared with another DELTEX family member DTX2. DTX3L shows preference for the 3’-terminal adenosine over other nucleotides. In addition, we demonstrate that ubiquitylation of nucleic acids is reversible by DUBs such as USP2 and SARS-CoV-2 PLpro. Overall, our study provides evidence for reversible ubiquitylation of nucleic acids *in vitro* and discusses its potential functional implications.

## INTRODUCTION

Regulating protein function by adding ubiquitin (Ub), known as ubiquitylation or ubiquitination, is widely utilized by eukaryotic cells. Ubiquitylation is involved in nearly all aspects of cellular activities, ranging from protein degradation, which was the first function of ubiquitylation to be discovered, to immune signaling, DNA damage response, receptor trafficking and many more (Oh *et al*, 2018; Zheng & Shabek, 2017). Ubiquitylation is the sequential transfer of Ub to the ε-amino group of a lysine residue on the substrate by a Ub-activating enzyme (E1), a Ub-conjugating enzyme (E2) and a Ub ligase (E3), resulting in the formation of an iso-peptide bond between Ub C-terminal glycine and the acceptor lysine residue (Hershko & Ciechanover, 1998; Komander & Rape, 2012). In addition, some E3s catalyze Ub transfer to the hydroxyl groups of threonine and serine residues or a thiol group of a cysteine residue, forming oxyester or thioester bond, respectively (Cadwell & Coscoy, 2005; Gao *et al*, 2021; Kelsall *et al*, 2019; Pao *et al*, 2018; Wang *et al*, 2009; Wang *et al*, 2007). Ub modification is highly reversible and detached by deubiquitinases (DUBs), with approximately a hundred of these enzymes encoded in the human genome (Mevissen & Komander, 2017).

Since ubiquitylation was first discovered five decades ago, tens of thousands of ubiquitylation sites on a large number of proteins have been identified, indicating that most proteins are ubiquitylated spatiotemporally in cells. It has been generally assumed that the substrates of ubiquitylation are solely limited to proteins. However, several recent studies changed this view, by demonstrating that Ub can be covalently attached to non-proteinaceous substrates, such as lipopolysaccharides (LPS) (Otten *et al*, 2021), phosphatidylethanolamine (PE) (Sakamaki *et al*, 2022), or glucosaccharides (Kelsall *et al*, 2022). Furthermore, Ub can also be attached on another modification called ADP-ribosylation (ADPr) (Zhu *et al*, 2023; Zhu *et al*, 2022). ADP-ribosylation is a chemical modification of proteins and nucleic acids involving the addition of one or more ADPr moieties. ADPr is transferred from nicotinamide adenine dinucleotide (NAD^+^) onto the targets by ADP-ribosyltransferases (ARTs) including the best-studied PARPs (Groslambert et al, 2021; Pascal, 2018; Suskiewicz et al, 2023b). Hybrid ADPr-Ub modification is efficiently synthesized *in vitro* by the DELTEX family of E3s on both proteins and nucleic acids substrates (Zhu *et al*., 2023; Zhu *et al*., 2022), but its physiological relevance is not clear yet.

DELTEX-family E3 ligases have been suggested to be involved in many pathways, for example, Notch signaling, DNA damage repair, innate immune response and cancer progression, and have attracted significant attention over the last decade, but the exact mechanisms and physiological consequences have been elusive (Wang et al, 2021). DELTEX family in human is composed of five members, namely DTX1, DTX2, DTX3, DTX4 and DTX3L (Takeyama et al, 2003; Wang et al., 2021). Different family members have distinct N-terminal domains, either WWE domains (in DTX1, DTX2 and DTX4), or KH domain(s) (in DTX3 and DTX3L)(Figure EV8) (Zhu et al., 2023). WWE domains in proteins are known to bind poly(ADP-ribose) chains (DaRosa *et al*, 2015) whereas KH domains are known for binding to single stranded nucleic acids (Nicastro *et al*, 2015a; Suskiewicz *et al*, 2023a; Valverde *et al*, 2008). In addition, DTX3L contains an RNA recognition motif (RRM) domain preceding the KH domains. In contrast to the varied N termini, DELTEX E3s share a characteristic C-terminal tandem RING-DTC domains (Chatrin et al, 2020), where the RING domain acts as an E3 Ub ligase and the DTC domain has been demonstrated to bind NAD^+^ and ADPr through its conserved pocket (Chatrin et al., 2020). Like other RING-type E3s, DELTEX RING domains don’t determine the specificity of Ub acceptors (a lysine amino group or a hydroxyl group), which is controlled by the E2s instead (Wenzel et al, 2011). However, DELTEX E3s evolved to have an accompanying DTC domain adjacent to RING domain, which bind NAD^+^ or ADPr and provide two catalytic residues to enable NAD^+^ or ADPr ubiquitylation on their 3’ hydroxyl groups of the adenine-proximal ribose (Zhu et al., 2022). Mechanistically, DELTEX E3s recruit E2∼Ub conjugate and one NAD^+^ or ADPr molecule using the RING and DTC domains, respectively. Next, the thioester bond between E2 and Ub is juxtaposed to the 3’ hydroxyl group of NAD^+^ or ADPr proximal ribose due to the flexible linker between RING domain and DTC domain (Zhu et al., 2022). Because the hydroxyl moiety is a weak nucleophile, DTC domain contributes one histidine residue and one glutamate residue to apparently deprotonate and thus encourage the 3’ hydroxyl group to attack the E2∼Ub conjugate to accomplish ADPr ubiquitylation.

Given the similarity between ADPr and nucleic acids, which are composed of the same constituents: nucleobases (adenine), ribose sugars and phosphates, we became intrigued by the possibility of directly ubiquitylating nucleic acids. Indeed, in this study, we demonstrate that DTX3L, representing KH domain-containing DELTEX E3s, ubiquitylates nucleic acids *in vitro*. The modification occurs primarily on the 3’-terminal adenosine nucleotide, likely targeting the 3’ hydroxyl group of the ribose sugar. Ubiquitylation of nucleic acids on the 3’ adenosine nucleotide protects them from degradation by 3’→5’ nucleases. Lastly, we show the reversibility of the DTX3L-mediated nucleic acids ubiquitylation by some DUBs including USP2 and SARS-CoV-2 PLpro.

## RESULTS

### DTX3L-RD ubiquitylates nucleic acids carrying the 3’ adenosine nucleotide

In our previous studies, we showed that the tandem RING-DTC (RD) domains of DELTEX family E3s are capable of ubiquitylating ADPr on the 3’ hydroxyl group of the adenine-proximal ribose (Zhu *et al*., 2023; Zhu *et al*., 2022). Specifically, the DTC domain first accommodates ADPr molecule to position the 3’ hydroxyl group of ADPr proximal ribose close to the E2∼Ub conjugate bound by the RING domain. The DTC domain appears to then utilize its two crucial catalytic residues to deprotonate the 3’ hydroxyl group, thus facilitating ADPr ubiquitylation. The available experimental structures show that the DTC domain uses the same conserved pocket to bind either ADPr or NAD^+^, and can facilitate Ub transfer to both (Figure 1A) (Ahmed *et al*, 2020; Chatrin *et al*., 2020; Zhu *et al*., 2023). Both ADPr and NAD^+^ contain an AMP moiety (Figure EV1), and it is this part that becomes ubiquitylated on the 3’ hydroxyl. By analyzing the ADPr/DTX2-RD and NAD^+^/DTX1-RD structures, we found in both complexes the shared AMP part of ADPr and NAD^+^ inserted into a deep pocket of DTC domains (Figure 1A). The AMP moiety has a highly similar conformation in the two structures and makes close contacts with the neighboring amino residues, making it distinguishable from the distal ribose of ADPr/NAD^+^, which protrudes out of the binding pocket, showing fewer contacts with the DTC domain. This finding prompted us to test whether AMP itself is also a substrate for ubiquitylation by DELTEX E3s. To test this, we utilized high-performance liquid chromatography coupled to mass spectrometry (HPLC-MS) to analyse the ubiquitylation reaction of DTX3L-RD with AMP, which shows that Ub is 100% converted into Ub-AMP (Figure EV2). This suggests that AMP is a substrate for ubiquitylation by DELTEX E3s, and its efficiency of ubiquitylation is comparable to that of ADPr or NAD^+^, both of which made more than 90% of starting Ub are conjugated to ADPr or NAD^+^ under the same conditions, as shown by HPLC-MS (Figure EV3) (Zhu *et al*., 2022).

**Figure 1.**
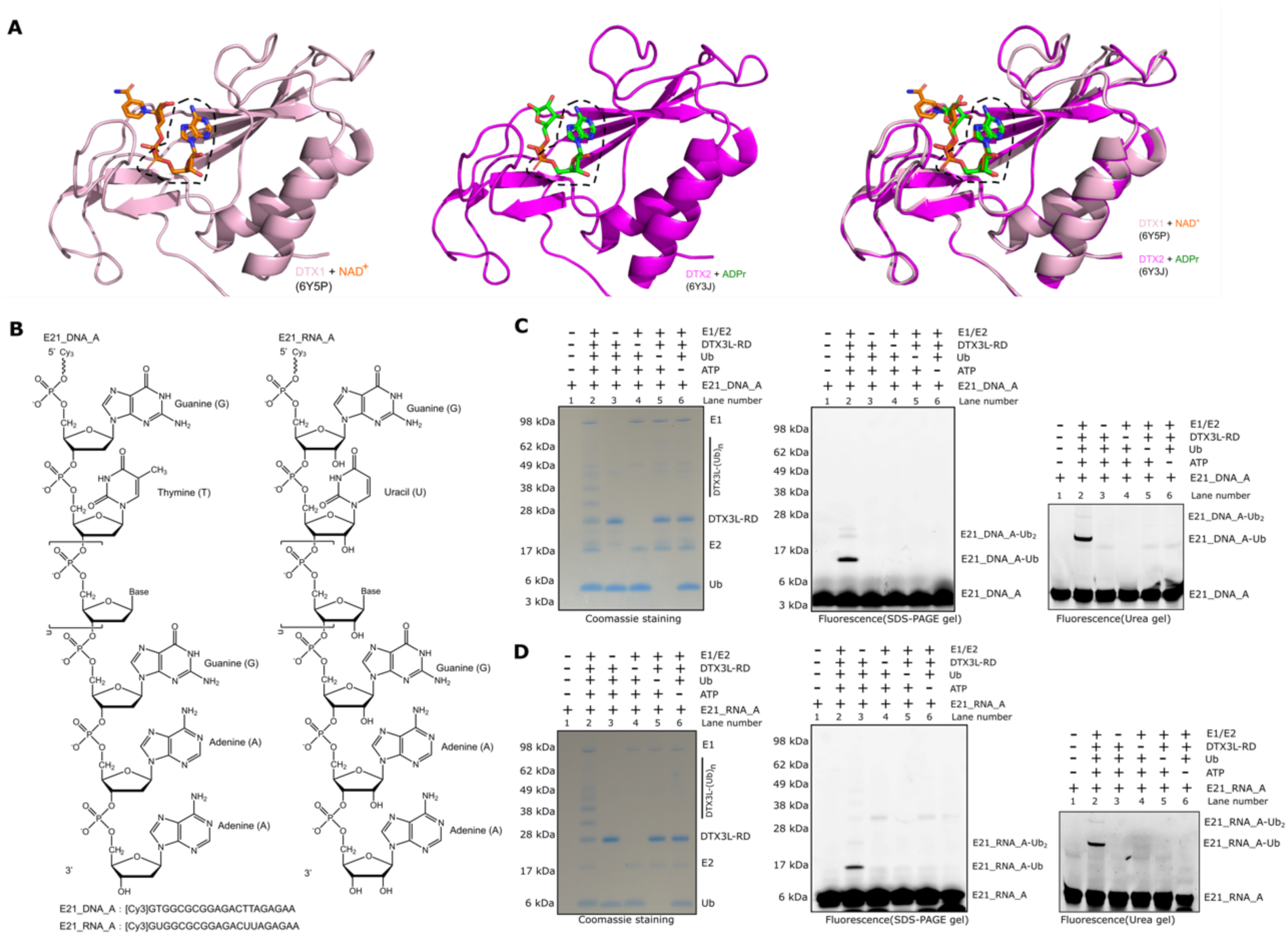
Biochemical characterization of DTX3L-RD-catalysed nucleic acids ubiquitylation. (A) Structural analysis showing that the shared AMP moiety in NAD^+^ and ADPr inserts deeply in the binding pockets of DTC domains from DTX1 or DTX2. The AMP moiety is indicated by black dashed box. (B) Chemical structures of the E21_DNA_A and E21_RNA_A used in this study. (C) Biochemical reconstitution of E21_DNA_A ubiquitylation. E21_DNA_A-Ub was obtained by incubation of DTX3L-RD and E1, E2, ATP and Ub. Omitting any of these components blocked the E21_DNA_A ubiquitylation. The reactions were divided into two parts. One part was analysed on an SDS-PAGE gel and visualized by first Molecular Imager PharosFX system (BioRad) and then coomassie staining. Another part was loaded on a pre-run 20% denaturing urea PAGE gel. The gels were run at 6 W/gel and following visualization using the Molecular Imager PharosFX system (BioRad). (D) As in (C), E21_RNA_A was used as substrate for the ubiquitylation reactions.

Considering that AMP or 2’ deoxy-AMP (dAMP) are building blocks for RNA or DNA, nucleic acids ending with adenosine nucleotide at the 3’ end will present an AMP/dAMP moiety with a free ribose 3’ hydroxyl, thus representing a potential ubiquitylation substrate for DELTEX E3s (Figure 1B). We wondered if RNA/DNA ending with (deoxy-)adenosine, could become directly ubiquitylated on their terminal riboses. To test this possibility, we designed a Cy3-labelled 21-nucleotide-long single-stranded DNA (ssDNA) with 3’ deoxy-adenosine nucleotide (E21_DNA_A) and selected DTX3L for the ubiquitylation assay, since DTX3L contains multiple single-stranded nucleic acids-binding domains (Zhu *et al*., 2023). We incubated DTX3L-RD with E21_DNA_A and Ub-processing components (E1, E2, Ub and ATP) and resolved the reaction mixtures on both SDS-PAGE and 20% TBE-Urea gels to visualize the potential nucleic acids-Ub adducts. As expected, in the presence of all ubiquitylation components, an upward-shifted band appeared, indicating that E21_DNA_A became ubiquitylated (Figure 1C, lane 2). However, the reactions omitting any ubiquitylation component did not show any higher band (Figure 1C, lane 3-6), which is consistent with what was observed for ADPr ubiquitylation (Zhu *et al*., 2023; Zhu *et al*., 2022). Similarly, we then used an ssRNA substrate that has the same sequence as E21_DNA_A (E21_RNA_A), and showed that E21_RNA_A was also ubiquitylated by DTX3L-RD (Figure 1D, lane 2). Full-length DTX3L (DTX3L fl) appears to be more efficient than DTX3L-RD in catalysing E21_DNA_A ubiquitylation, possibly owing to enhanced substrate recruitment through multiple nucleic acids-binding domain (Zhu *et al*., 2023)(Figure EV4). However, since the minimum catalytic RING-DTC (RD) fragment is proficient enough and easier to produce, this fragment is used throughout the study.

Next, we wanted to figure out on which chemical moiety within nucleic acids Ub is attached. Considering the chemistry of the ubiquitylation reaction, with E2∼Ub acting as an electrophile, we focussed on nucleophilic moieties that could act as Ub acceptors. Depending on whether DNA or RNA is used, one or two hydroxyl group(s) on the 3’-terminal adenosine are available, in addition to several amine groups on terminal or internal bases (Figure 1B). However, considering the similarity between adenosine nucleotide and ADPr, the 3’ hydroxyl group of the 3’-terminal ribose appears the most likely candidate (Zhu *et al*., 2023). Of note, the 3’ hydroxyl group, unlike the 2’ hydroxyl group, is shared between DNA and RNA molecules, both of which were efficiently ubiquitylated above. We used phosphorylation to block the 3’ hydroxyl group of the terminal ribose, and tested if it affected the Ub modification. We conducted ubiquitylation reactions using DTX3L-RD and E21_DNA_A as well as its 3’ phosphorylated form, E21_DNA_A_3P. In contrast to E21_DNA_A, which was ubiquitylated, E21_DNA_A_3P ubiquitylation was greatly weakened, suggesting that the terminal 3’ hydroxyl group is the likely Ub acceptor site (Figure 2A). The weak remaining ubiquitylation of the phosphorylated DNA might be due to incomplete phosphorylation. Moreover, considering that the ester bond between the Gly^76^ residue of Ub and the 3’ hydroxyl group of terminal adenosine nucleotide should be sensitive to NH_2_OH treatment (Zhu *et al*., 2023; Zhu *et al*., 2022), we used NH_2_OH to see if it can reverse the ubiquitylation of E21_DNA_A and E21_RNA_A. Our result showed that NH_2_OH completely removed the ubiquitylation, speaking against the possibility of amide group-linked ubiquitylation, which is resistant to NH_2_OH (Figure 2B, Figure EV5A). Consistent with this, ubiquitylation of E21_DNA_A and E21_RNA_A was abolished upon mutating catalytic histidine and glutamate residues (H707A and E733R) present in the DTC domain of DTX3L-RD (Figure 2C, Figure EV5B), which are required for ADPr ubiquitylation on 3’ hydroxyl but not canonical lysine ubiquitylation (Zhu *et al*., 2023; Zhu *et al*., 2022).

**Figure 2.**
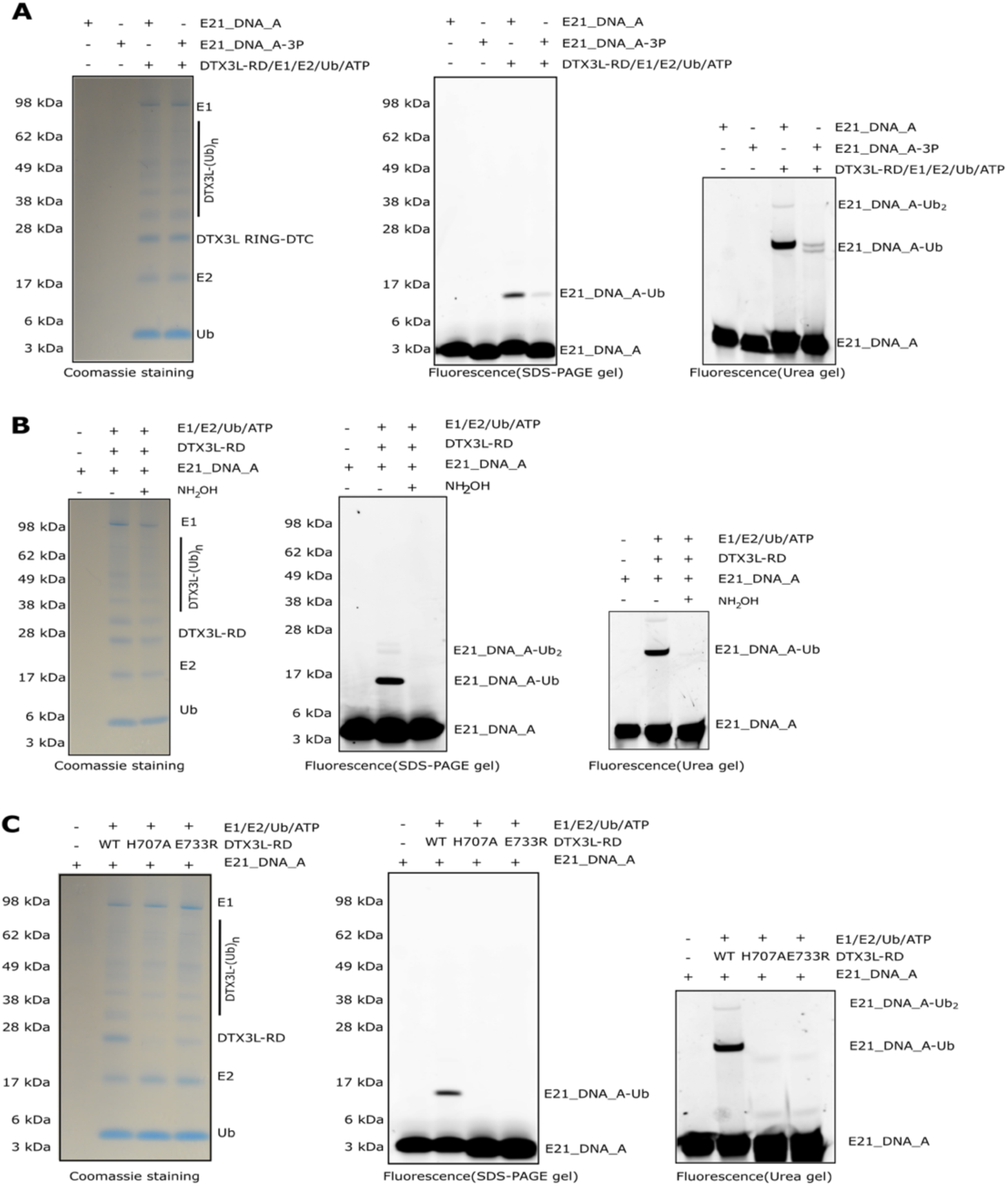
DTX3L-RD attaches Ub onto the 3’ hydroxyl group of terminal adenosine in nucleic acids. (A) 3’ phosphorylation markedly reduced the ubiquitylation of DNA by DTX3L-RD. E21_DNA_A and its 3’ phosphorylation form: E21_DNA_A-3P were incubated with E1, E2, ATP and Ub. The reactions were analysed on an SDS-PAGE gel and urea PAGE gel and processed as described before. (B) NH_2_OH reverses DTX3L-RD-catalysed E21_DNA_A ubiquitylation. NH_2_OH cleaves the ester bond between the carbonyl group of Gly^76^ of Ub and the 3’ hydroxyl group of the A of nucleic acids. (C) DTX3L-RD ADPr ubiquitylation inactive mutants failed to produce upshift bands that correspond to ubiquitylation of DNA, indicating that Ub is attached to 3’ hydroxyl group.

Overall, these results suggest that the observed ubiquitylation of nucleic acids happens on the 3’-terminal adenosine nucleotide, likely through its 3’ hydroxyl group.

### DTX3L-RD shows preference for 3’-terminal adenosine nucleotide over other nucleotides in nucleic acids

Since we showed that DTX3L-RD could ubiquitylate the 3’ adenosine (A) of nucleic acids, we therefore wondered whether other nucleotides at the 3’ end could also be ubiquitylated. According to their chemical structures, the purine GMP resembles AMP with a two-ring structure, while pyrimidines CMP and TMP have a different one-ring structure (Figure 3A). To test our idea, we first utilized HPLC-MS to analyse the ubiquitylation reaction of DTX3L-RD with free nucleotides including GMP, CMP and TMP (Figure EV6-S8), using AMP and ADPr as the control (Figure EV2 and EV3). Interestingly, we detected the masses that are consistent with the molecular weights of Ub-GMP, Ub-CMP and Ub-TMP, but the efficiency of ubiquitylation, judged by the percentage of nucleotides that became modified, varied and were generally lower than for Ub-AMP which was the only nucleotide that can be quantitatively modified (Figure 3B) indicating that AMP is preferred by DTX3L-RD. In all cases, the only Ub-containing products detected were the starting Ub and Ub-NMP (N=A, T, C or G), together with an Ub-DTT adduct in the case of the poor substrates GMP, TMP, and CMP. This adduct is likely formed upon direct nucleophilic attack of DTT to the E2∼Ub conjugate, through a trans-thioesterification process.

**Figure 3.**
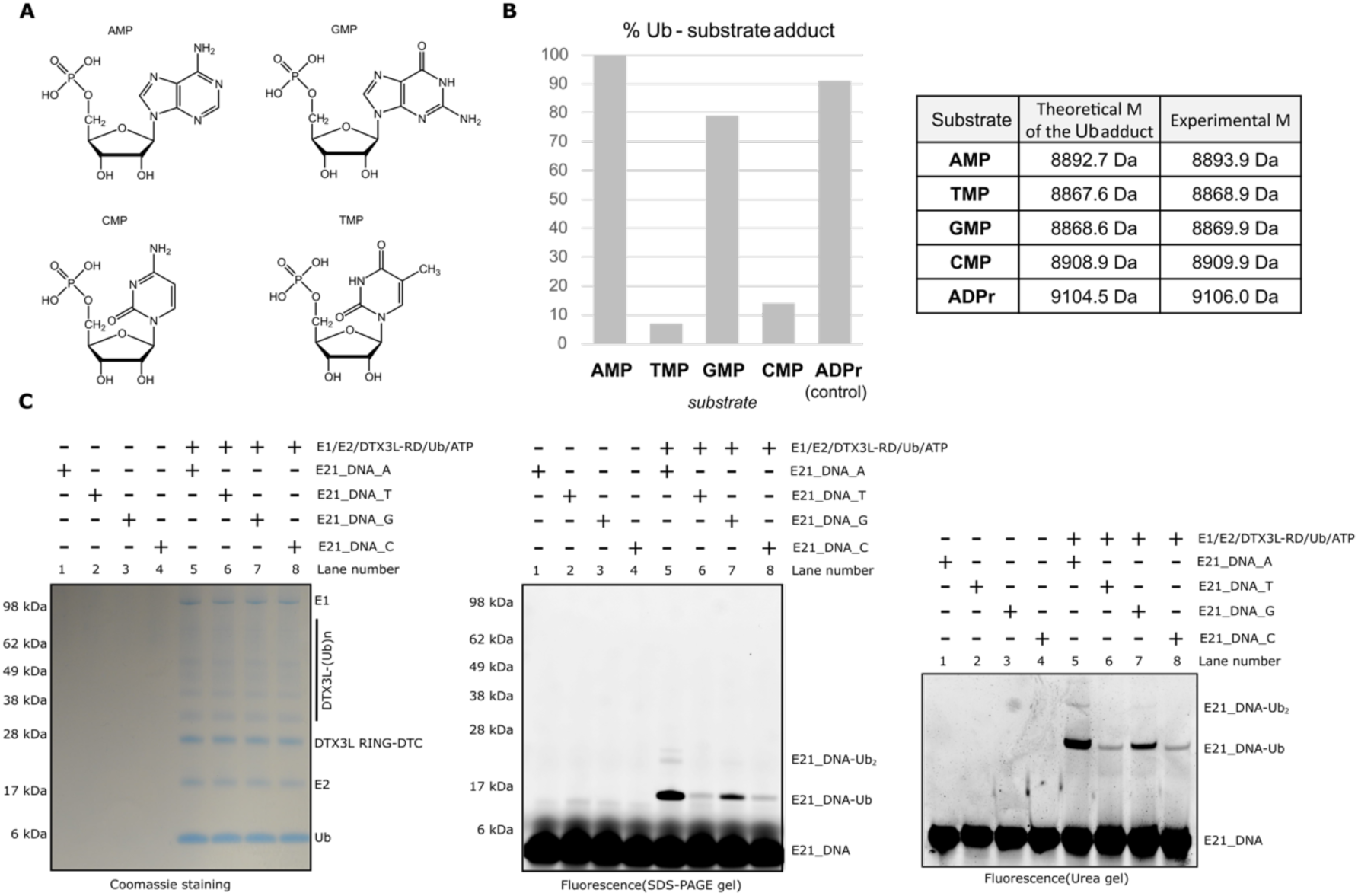
DTX3L-RD preferably ubiquitylates 3’ adenosine base of nucleic acids. (A) Chemical structures of different nucleotides. (B) HPLC-MS-based identification of the products of DTX3L-RD-catalysed ubiquitylation reactions performed with indicated nucleotides. Detected average masses and theoretical ones are provided. Ub was used at a 25 µM concentration, and all substrates at 12 mM (48 molar equivalents). (C) DTX3L-RD ubiquitylates 3’ A, G, T and C of nucleic acids. E21_DNA_A-Ub was obtained by incubation of DTX3L-RD and E1, E2, ATP and Ub. The reactions were divided into two parts. One part was analysed on an SDS-PAGE gel and visualized by first florescence and then coomassie staining. Another part was loaded on a pre-run 20% denaturing urea PAGE gel. The gels were run at 6 W/gel and following visualization using the Molecular Imager PharosFX system (BioRad).

Next, we wanted to know whether DTX3L-RD could ubiquitylate nucleic acids with 3’ G, T or C ends and whether the efficiency of the potential ubiquitylation shows a similar trend to that observed with free nucleotide monophosphates. We redesigned the E21_DNA_A to have different 3’ ends, namely, T, C and G ends, and tested these in our ubiquitylation assay. Our results showed the most abundant ubiquitylation for E21_DNA_A, followed by E21_DNA_G, while only a small fraction of E21_DNA_T and E21_DNA_C were ubiquitylated (Figure 3C). This data is consistent with our MS data obtained with free nucleotides.

Taken together, our results suggest that DTX3L preferably ubiquitylates 3’-terminal adenosine of nucleic acids over other 3’-terminal nucleotides. Additionally, the strict dependence of the reaction efficiency on the nature of the 3’-terminal nucleotide further supports the notion that the modification takes place on that nucleotide.

### DTX3L possesses nucleic acids ubiquitylation activity while DTX2 does not

Based on their N-terminal domains, DELTEX family E3s can be divided into two sub-classes (Figure EV9A): (1) WWE domain-containing DELTEX E3s: DTX1, DTX2 and DTX4, all of which possess two WWE domains (Wang *et al*., 2021; Zweifel *et al*, 2005); and (2) KH domain-containing DELTEX E3s: DTX3 and DTX3L, where DTX3 contains one KH domain while DTX3L has five KH domains (Zhu *et al*., 2023). Considering that the WWE domain typically binds the poly(ADP-ribose) chain (DaRosa *et al*., 2015; Kang *et al*, 2011; Wang *et al*, 2012), while the KH domain tends to bind single-stranded nucleic acids (Nicastro *et al*, 2015b; Valverde *et al*., 2008), we speculated that this might reflect a functional difference, with KH domain-containing DELTEXes having a nucleic acids-related function. Our results so far indicated that DTX3L-RD ubiquitylates nucleic acids, preferably on its 3’-terminal adenosine. We wondered whether DTX2, a WWE domain-containing DELTEX, is capable of ubiquitylating nucleic acids. To test this, we performed ubiquitylation reaction using DTX2-RD while DTX3L-RD was used as the control. Surprisingly, DTX2-RD did not show any ubiquitylation activity on E21_DNA_A, even at 16 µM enzyme concentration, whilst DTX3L-RD ubiquitylated E21_DNA_A at 1 and 4 µM concentration (Figure EV9B). This suggests that DTX3L exhibits nucleic acids ubiquitylation activity, whereas DTX2 does not. Since, in the above experiment, RD fragments of DTX3L and DTX2 were used, the difference does not come from the presence or absence of KH domains, but rather is inherent to the RD fragment, which apparently evolved differently in the two sub-classes of DELTEX E3 ligases.

### Ubiquitylation of nucleic acids prevents degradation and is reversible

Chemical modifications on nucleic acids plays important roles in their function including influencing their stability. For example, eukaryotic mRNAs undergo co-transcriptional modification through the addition of a 7-methylguanosine cap (m7G), which shields mature mRNAs from degradation by 5’→3’ exonucleases (Furuichi *et al*, 1977; Shatkin, 1976). In contrast, NAD^+^ capping at the 5’ end of RNA has been observed to promote degradation (Jiao *et al*, 2017; Yu *et al*, 2021). Recent studies have reported ADPr as another capping mechanism of the 5’ end of RNA, which protects them from degradation by nucleases, thus improving their stability (Munnur *et al*, 2019). Prompted by these findings, we investigated whether ubiquitylation of nucleic acids at their 3’ end influence their stability. We first used DTX3L-RD and E21_RNA_A to generate ubiquitylated E21_RNA_A, then treated the reaction mix with 3’→5’ exonuclease T (Exo T). We observed that unmodified E21_RNA_A was completely degraded, while the ubiquitylated E21_RNA_A was resistant to Exo T treatment (Figure 4A), suggesting the protective role of ubiquitylation in this specific manner.

**Figure 4.**
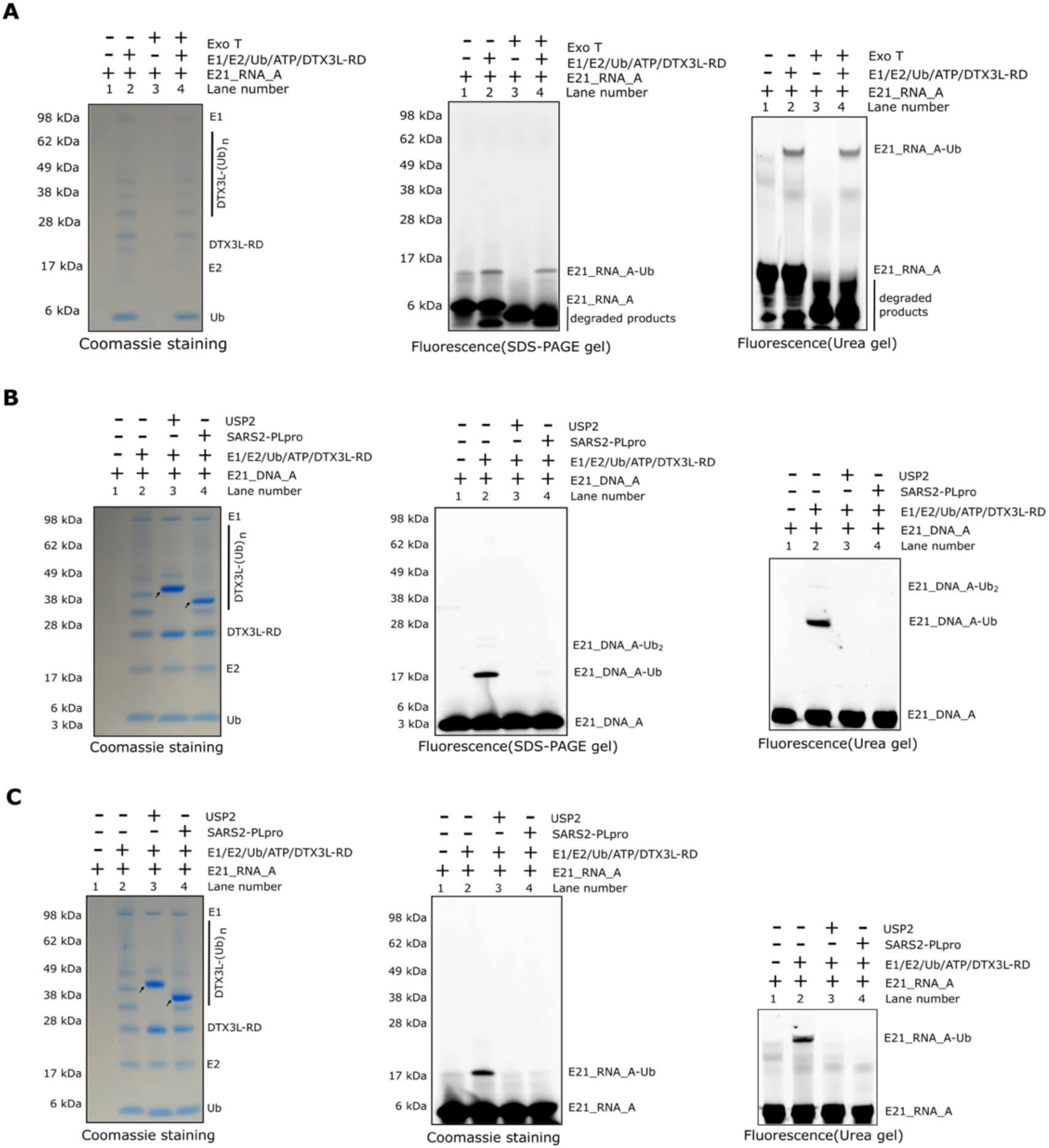
Ubiquitylation of nucleic acids protect them from 3’→5’ nuclease attack and is a reversible process. (A) The ubiquitylation of E21_RNA_A prevents cleavage by the 3’-5’ nuclease Exo T. Ubiquitylated E21_RNA_A and unmodified E21_RNA_A were treated with Exo T and the reactions were resolved via SDS-PAGE gel and Urea gel. RNA and RNA-Ub adduct were visualised by the Molecular Imager PharosFX system (BioRad). (B) Hydrolysis of ubiquitylated nucleic acids. Following the ubiquitylation of E21_DNA_A with DTX3L-RD, the indicated DUBs were added and further incubated. The reactions were analysed and visualized as described earlier. The arrows indicate various hydrolases. (C) As in (B), ubiquitylated E21_RNA_A was used as substrates for hydrolysis. The reactions were analysed and visualized as described earlier.

Following the characterization of nucleic acids ubiquitylation by DTX3L, we next wanted to test whether the Ub modification on nucleic acids is a reversible process. Modifications of proteins and nucleic acids can typically be reversed by the action of so-called eraser enzymes (DUBs in the case of ubiquitylation), which can remove the added chemical groups. We first generated ubiquitylated E21_DNA_A using DTX3L-RD and then treated the reaction mix with USP2, an Ub-substrates linkage-nonspecific DUB (Amerik & Hochstrasser, 2004; Renatus *et al*, 2006). Notably, USP2 removed the Ub modification from E21_DNA_A (Figure 4B). Additionally, we also included SARS2-CoV-2 PLpro, since SARS2 virus infection causes a strong induction of DTX3L, which has been reported to function as an antiviral protein (Heer *et al*, 2020). Our results showed that SARS-CoV-2 PLpro reversed ubiquitylation of E21_DNA_A completely (Figure 4B). In an analogous set of experiments, we reproduced the same observations with the ssRNA E21_RNA_A (Figure 4C).

Taken together, our data show that ubiquitylation of nucleic acids on the 3’ adenosine can function as a protective mechanism against 3’→5’ nucleases such as ExoT *in vitro*, suggesting the potential protective role in cells. Furthermore, we demonstrated that nucleic acids ubiquitylation can be enzymatically reversed by USP2 and SARS-CoV-2 PLpro.

## DISCUSSION

Ubiquitylation plays a pivotal role in nearly all cellular processes, including protein homeostasis, immune signaling, and DNA damage response (Hershko & Ciechanover, 1998; Komander & Rape, 2012). Alterations in ubiquitylation can detrimentally affect the regulation of key signaling pathways, exacerbating cellular dysfunction, and contributing to pathological conditions. However, our understanding of ubiquitylation remains limited, particularly regarding the diverse types of ubiquitylation substrates. Since it was discovered, ubiquitylation has been considered to be exclusive to proteins, as all reported acceptors of ubiquitin were lysine, threonine, or serine residues in protein substrates. However, in 2021, the discovery of lipopolysaccharides (LPS) brought attention to non-proteinaceous substrates of ubiquitylation, opening up a new frontier in the Ub field (Otten *et al*., 2021). Subsequently, glucosaccharides (sugar), phosphatidylethanolamine (lipids), and ADPr were identified as ubiquitylated molecules (Kelsall, 2022; Sakamaki & Mizushima, 2023; Zhu *et al*., 2022). Nevertheless, a lingering question has remained: are there other non-proteinaceous substrates of ubiquitylation?

In advancing the field, we previously demonstrated that ADPr is a non-proteinaceous substrate for DELTEX E3s-mediated ubiquitylation, with the 3’ hydroxyl group of the adenine-proximal ribose of ADPr serving as the Ub acceptor. Interestingly, the ADPr that is attached to a peptide, a protein, or a nucleic acid can also be ubiquitylated by DELTEX E3s, thus allowing indirect ubiquitylation of various substrates (Zhu *et al*., 2023; Zhu *et al*., 2022). Considering the similarity between ADPr and nucleotides, the building blocks of nucleic acids, we now investigated whether DELTEX E3 ligases could directly ubiquitylate nucleotides and nucleic acids. Indeed, in the current study, we show that DTX3L ubiquitylates free nucleotide monophosphates as well as nucleic acids, with a preference for those carrying 3’-terminal adenosine nucleotides. Blocking the 3’ ribose hydroxyl group of the terminal adenosine prevents the modification, in line with the hypothesis – based on the mechanism of ADPr ubiquitylation – that the 3’ hydroxyl group serves as the Ub target. Our study establishes nucleic acids as a novel type of ubiquitylation substrate.

Do all members of the DELTEX family possess the ability to ubiquitylate nucleic acids? We first tested DTX3L, which, in its full-length version, harbors single-stranded nucleic acids-binding domains, and it could catalyse ubiquitylation of nucleic acids. In contrast, we did not observe nucleic acids ubiquitylation with DTX2-RD, even after using very large amount of the protein. This suggests that the small differences in the catalytic RD fragments (Figure EV9C) can lead to or exclude the nucleic acids ubiquitylation activity. In NAD^+^ or ADPr ubiquitylation, the catalysis is strictly coordinated by the RING domain, DTC domain, and their specific orientation via the flexible linker between them. A previous study showed that mixing isolated RING domain and DTC domain or changing the length or flexibility of the linker abolish ADPr ubiquitylation activity, indicating the requirement for precise coupling between the two domains (20). It is likely that nucleic acids ubiquitylation also follows a similar principle, with specific structural requirements within and possibly between the two domains. Considering the differences between DTX3L (KH domain-containing DELTEXes) and DTX2 (WWE domain-containing DELTEXes), we speculated that the slightly shorter RING domain and/or the lack of an AR insertion in DTC domain of DTX3L might affect its ability in ubiquitylating nucleic acids (Figure EV9C). It is likely that other unidentified elements may also be essential. Hence, a more detailed investigation is warranted for a comprehensive understanding of the difference between the two DELTEX sub-classes with respect to their ubiquitylation substrate specificity.

Covalent conjugation of proteins and nucleic acids could have regulatory consequences to both proteins and nucleic acids. A noteworthy recent study exemplifies this possibility, revealing that the bacteriophage T4 employs its ARTs to engage in an ‘RNAylation’ reaction (Wolfram-Schauerte *et al*, 2023). In this process, RNA chains are added to host *Escherichia coli* ribosomal proteins, strategically targeting the translational machinery and contributing to the bacteriophage’s pathogenicity. Although the traditional modifications of nucleic acids has predominantly involved the attachment of small chemical groups, such as the methyl (for example in the case of N6-methyladenosine, m6A), our investigation demonstrates that nucleic acids can undergo modification with a large, proteinaceous modifier (Ub). Furthermore, nucleic acids ubiquitylation can be reversed by some DUBs including USP2 and SARS-CoV-2 PLpro, indicating that this process is a reversible reaction. Despite the current lack of understanding regarding the physiological function of the uncovered process, our preliminary results suggested that DTX3L-catalyzed ubiquitylation on 3’-terminal adenosine end of nucleic acids protects it from degradation by 3’→5’ exonucleases *in vitro*. Given that the poly(A) tail is crucial for mRNA stability, transport, and translation, we propose that ubiquitylation of the poly(A) in mRNA may add another regulatory layer to these processes. Although our data only show the nucleic acids ubiquitylation activities of DTX3L *in vitro*, we speculate that these products might have *in vivo* implications: the protection of nucleic acids’ 3’ ends, blocking DNA end processing during DNA repair, influencing the nucleic acids’ stability, interactome, and translational processes. However, it is also possible that the nucleic acids ubiquitylation activity by DTX3L is a detrimental off-target activity of these DELTEX E3s, which – when happening in the cell – would need to be repaired. In such a scenario, DUBs such as USP2 would function as a repair enzyme by reversing the aberrant ubiquitylation of nucleic acids, analogously to the repair of DNA adenylates formed during abortive DNA ligation events by aprataxin (APTX) (Ahel *et al*, 2006). Similarly, it has been suggested that several PARPs mistakenly ADP-ribosylate DNA, thereby generating DNA lesions (adducts), which could be repaired by ADPr hydrolases (Munnur & Ahel, 2017).

In summary, our data show that the DTX3L E3 Ub ligase can ubiquitylate nucleic acids at the 3’ terminus and that this modification can be removed by USP2 and SARS-CoV-2 PLpro. This surprising discovery suggests that reversible ubiquitylation of nucleic acids can happen at least *in vitro,* but may also be a novel strategy utilized in cellular signaling in the context of some uncharacterized ubiquitylation systems from different organisms. Further research would be necessary to fully understand the functional significance of nucleic acids ubiquitylation.

## MATERIALS AND METHODS

### Plasmids and protein purification

WT and mutants of the RING-DTC domains of DTX3L (DTX3L-RD) and DTX2 (DTX2-RD) were expressed and purified as previously described (Zhu *et al*., 2023; Zhu *et al*., 2022). SARS-CoV-2 PLpro and USP2 were produced recombinantly before in our laboratory. Full-length DTX3L (DTX3L fl) was expressed and purified as previously described (Zhu *et al*., 2023; Zhu *et al*., 2022).

UBE1 (E-304-050), UBCH5A (E2-616-100), and recombinant Ub (U-100H-10M) were purchased from R&D Systems. Exo T nuclease (M0265S) was purchased from NEB.

### Oligonucleotide

Single-stranded (ss) DNA or RNA oligos used in this study were commercially ordered from Sigma-Aldrich. Oligonucleotides were dissolved to 100 μM stock in 20 mM HEPES–KOH (pH 7.6) and 50 mM KCl buffer.

Sequence of oligonucleotides used in this study (5′→3′):

5Cy3_E21_DNA_A [Cy3]GTGGCGCGGAGACTTAGAGAA

5Cy3_E21_DNA_T [Cy3]GTGGCGCGGAGACTTAGAGAT

5Cy3_E21_DNA_G [Cy3]GTGGCGCGGAGACTTAGAGAG

5Cy3_E21_DNA_C [Cy3]GTGGCGCGGAGACTTAGAGAC

5Cy3_E21_DNA_A_3’P [Cy3]GTGGCGCGGAGACTTAGAGAA[Phos]

5Cy3_E21_RNA_A [Cy3]GUGGCGCGGAGACUUAGAGAA

### Nucleic acids ubiquitylation assay

4 μM DTX3L RING-DTC was incubated with 0.5 μM UBE1, 2.5 μM UBCH5A, 10 μM Ub, and 1μM Cy3-labelled individual nucleic acids in 50 mM HEPES pH 7.5, 50 mM NaCl, 5 mM MgCl_2_, 1 mM DTT, and 1 mM ATP. After incubation at 37 °C for 1 h, reactions were split into two equal parts. One half was stopped by addition of 4X LDS sample buffer (Life Technologies) and analyzed by SDS-PAGE gel, which was first imaged using the Molecular Imager PharosFX system (BioRad) with laser excitation for Cy3 at 532 nm, then stained with Coomassie staining. Another half was stopped by addition of 2x TBE urea sample buffer (8 M urea, 20 µM EDTA pH 8.0, 20 µM Tris-HCl pH 7.5, and bromophenol blue) and loaded on a pre-run 20% denaturing urea PAGE gel. The gels were run at 6 W/gel and followed by Cy3 visualization using the Molecular Imager PharosFX system (BioRad).

For Figure EV4, 4 μM DTX3L-RD or 0.5 μM DTX3L fl was used.

### Nucleic acids ubiquitylation blocks Exo T’s activity

4μM DTX3L-RD was incubated with 0.5 μM UBE1, 2.5 μM UBCH5A, 10 μM Ub, and 1μM Cy3-labelled individual nucleic acids in 50 mM HEPES pH 7.5, 50 mM NaCl, 5 mM MgCl_2_, 1 mM DTT, and 1 mM ATP. After incubation at 37 °C for 1 h, 10 mM EDTA was used to stop the reactions and then 1U Exo T per reaction was added and incubated at 25 °C for 30 min. The samples were resolved using SDS-PAGE gel and Urea gel and visualized as described above.

### HPLC-MS analyses

HPLC-MS analyses were carried out on an Agilent 1260 Infinity HPLC system, coupled with an Agilent 6120 mass spectrometer [electrospray ionization (ESI) + mode]. The multiply charged envelope was deconvoluted using the charge deconvolution tool in Agilent OpenLab CDS ChemStation software to obtain the average [M] value.

### HPLC-MS monitoring of Ub-NMPs (N=A, T, G or C)

Ub-NMPs were generated by incubation of 12 mM individual NMP with 5 μM DTX3L RING-DTC, 0.5 μM UBE1, 2.5 μM UBCH5A, and 20 μM Ub in 50 mM HEPES (pH 7.5), 50 mM NaCl, 5 mM MgCl_2_, 0.5 mM DTT, and 2 mM ATP. Post incubation at 37°C for 2h, 10 μl reactions were mixed with 2 μl of 1% TFA. Then reactions were subjected to HPLC-MS analysis as previously described (Zhu *et al*., 2023; Zhu *et al*., 2022).

## AUTHOR CONTRIBUTIONS

K.Z., D.A. and I.A. conceived and designed the experiments. K.Z. and C.C. conducted the biochemical experiments. V.A. performed HPLC-MS analysis. K.Z. and C.C. performed structural analysis and prepared the figures. K.Z., M.J.S and C.C. wrote the original manuscript.

## ACKNOWLEDGEMENTS

We thank Zining Zhu for preparing reagents.

---

**Expanded View Figure 1.**
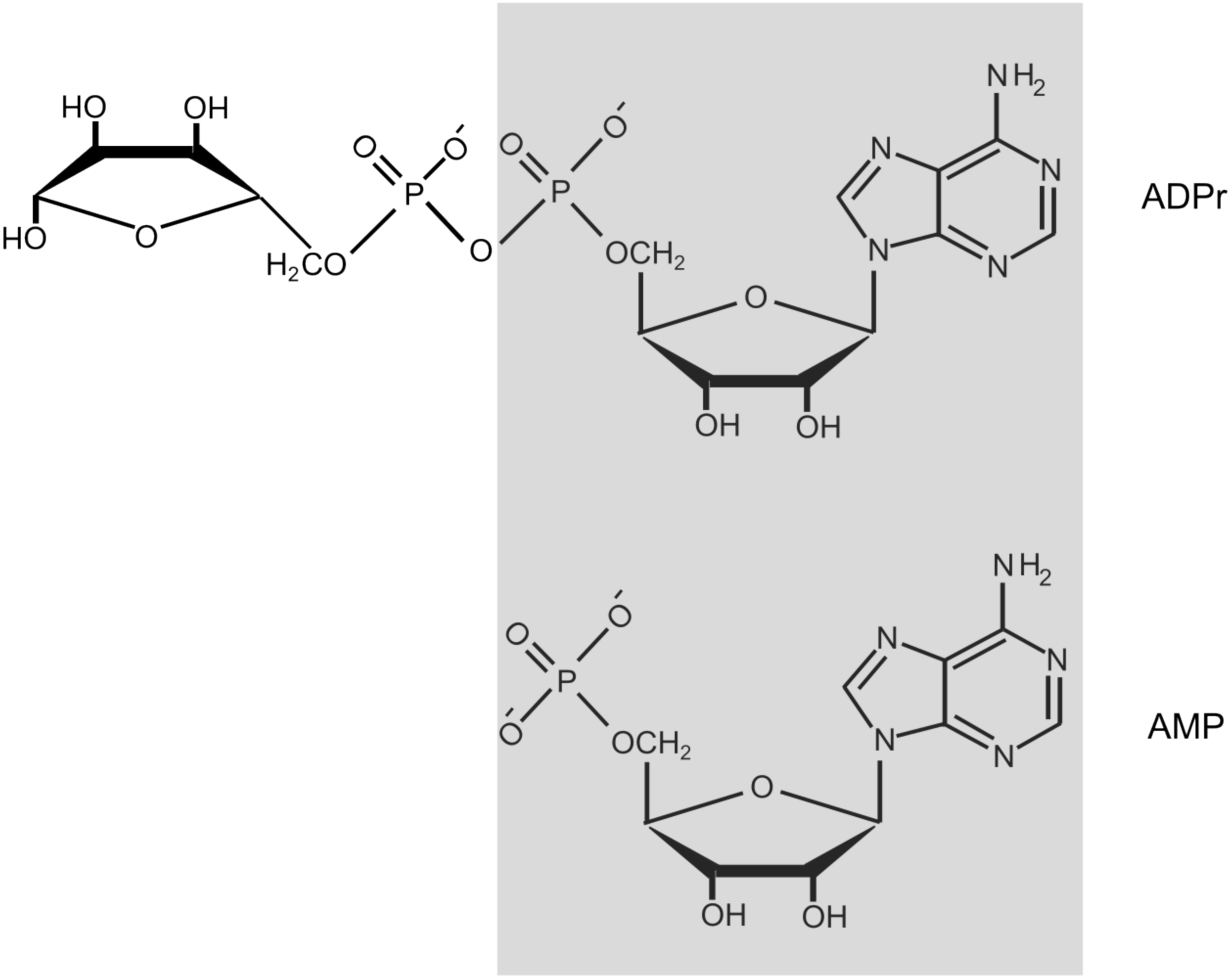
Chemical structures of AMP and ADPr. ADPr contains AMP core in its structure.

**Expanded View Figure 2.**
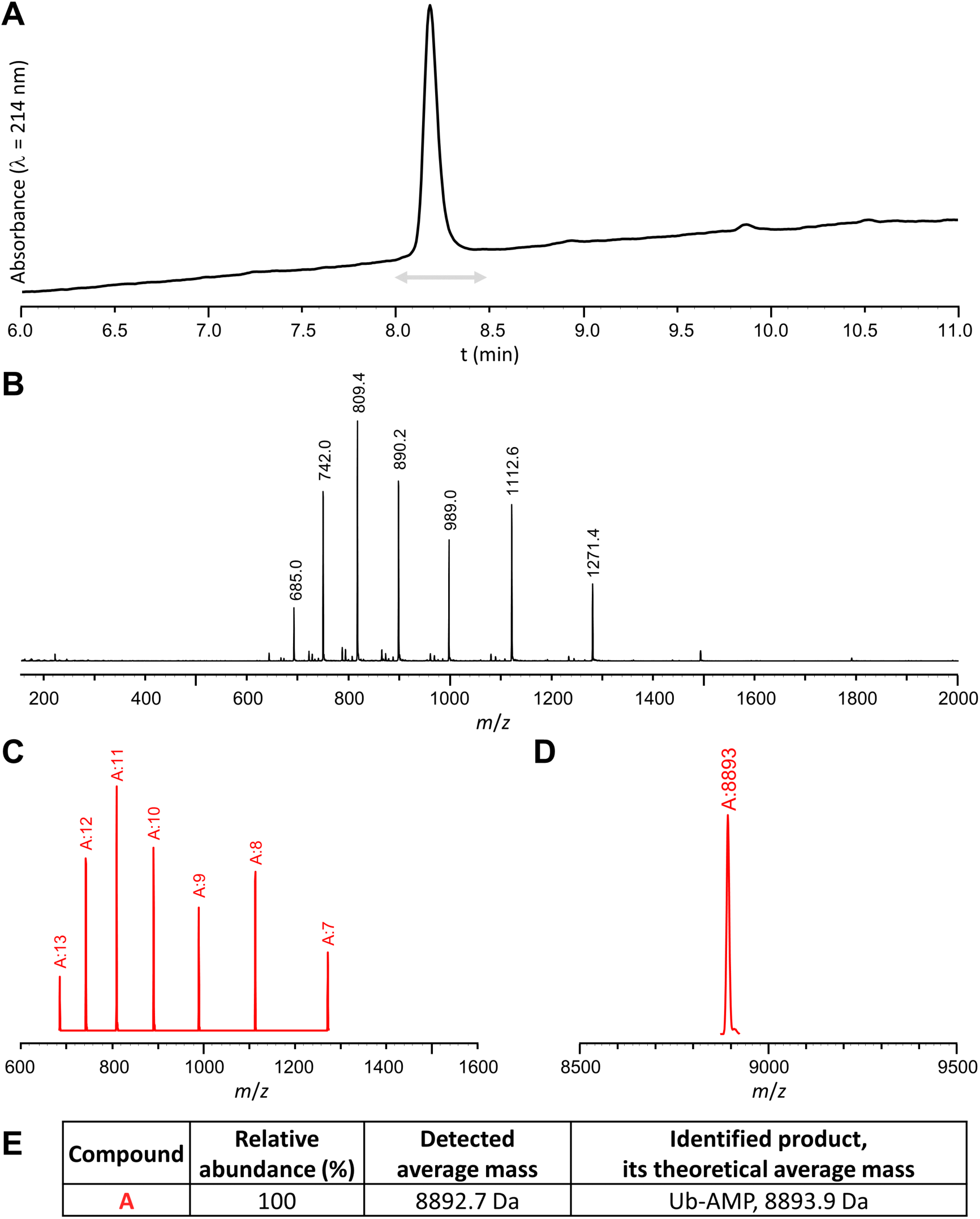
HPLC-MS analysis of the ubiquitylation mixture performed using DTX3L-RD and AMP. (A) HPLC chromatogram; (B) Experimental mass spectrum corresponding to the time window indicated as a grey arrow (sum of spectra); (C) Deconvoluted ions set, including charge state; (D) Deconvoluted spectrum; (E) Identified compounds.

**Expanded View Figure 3.**
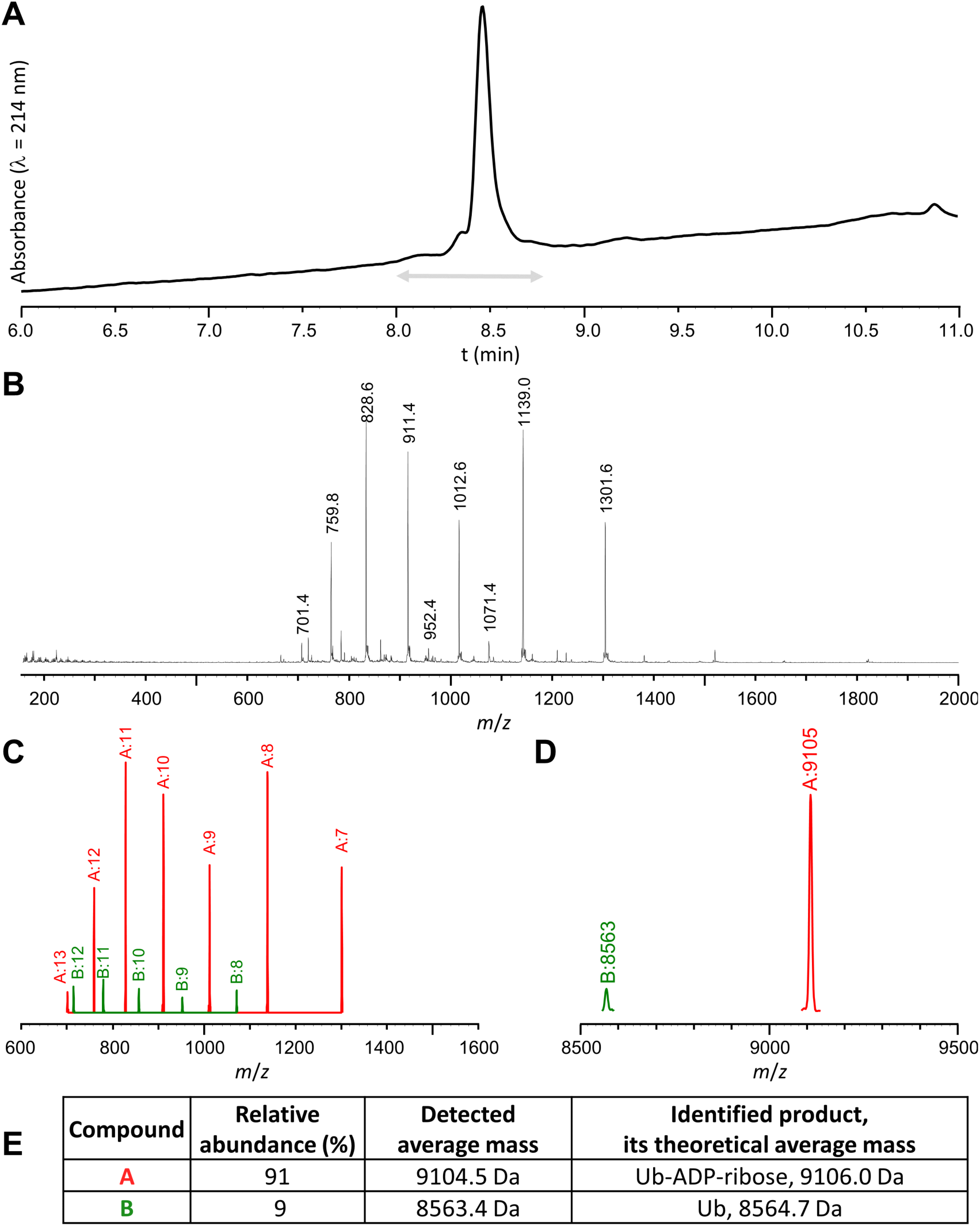
HPLC-MS analysis of the ubiquitylation mixture performed using DTX3L-RD and ADPr. (A) HPLC chromatogram; (B) Experimental mass spectrum corresponding to the time window indicated as a grey arrow (sum of spectra); (C) Deconvoluted ions set, including charge state; (D) Deconvoluted spectrum; (E) Identified compounds.

**Expanded View Figure 4.**
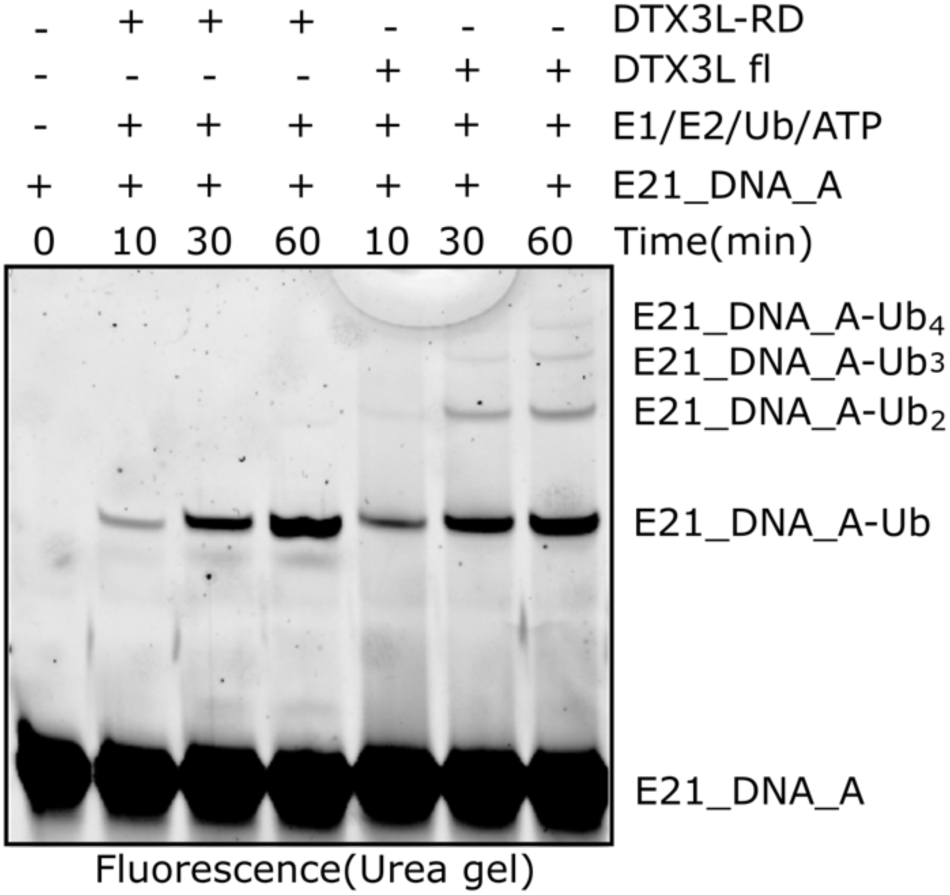
DTX3L catalysed nucleic acids ubiquitylation. E21_DNA_A ubiquitylation by DTX3L-RD and DTX3L fl, at indicated time points.

**Expanded View Figure 5.**
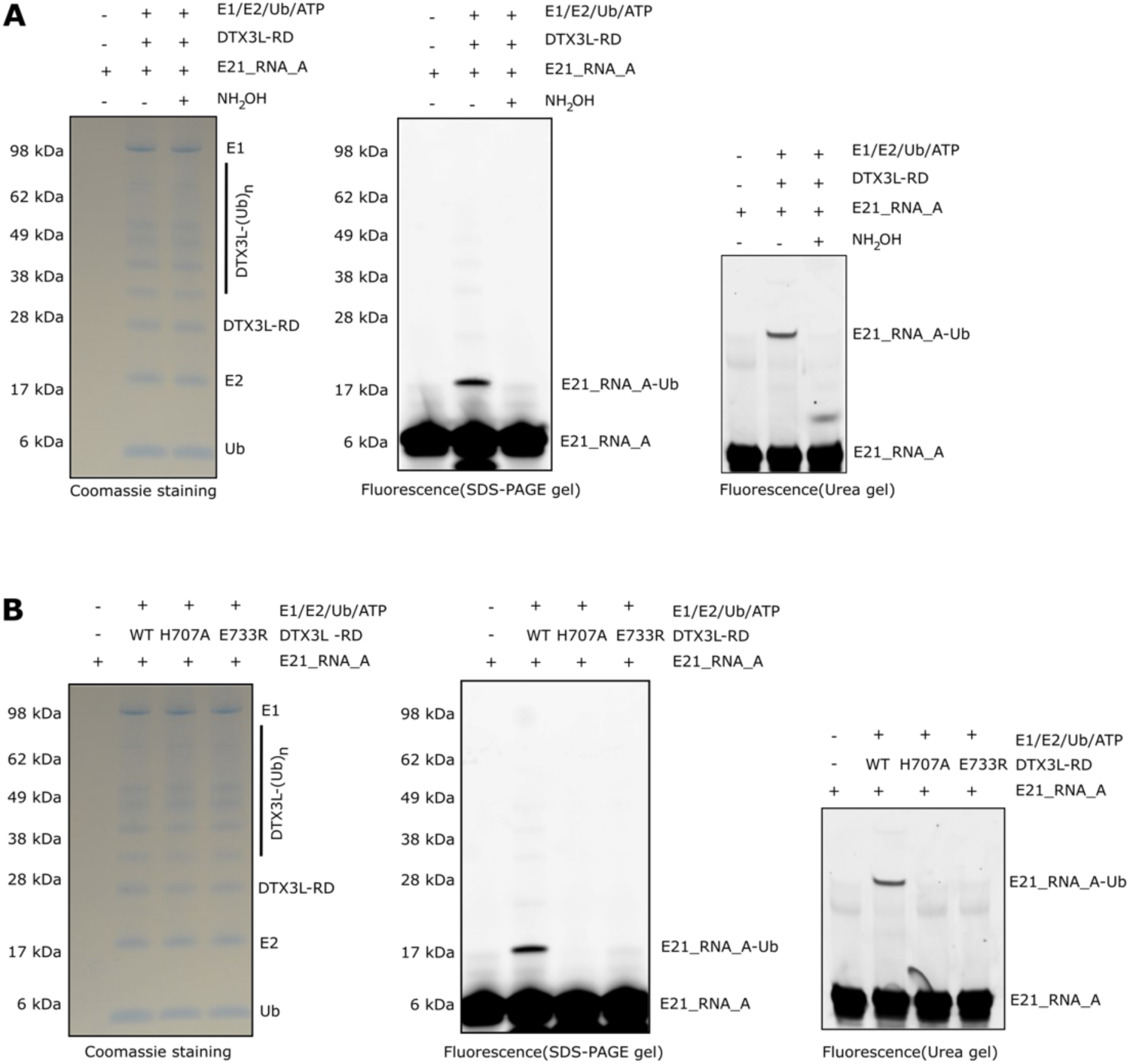
DTX3L-RD attaches Ub onto the 3’ hydroxyl group of terminal adenosine in RNA. (A) NH_2_OH reverses DTX3L-RD-catalysed nucleic acids ubiquitylation. NH_2_OH cleaves the ester bond between the carbonyl group of Gly^76^ of Ub and the 3’ hydroxyl group of the A of E21_RNA_A. (B) DTX3L-RD ADPr ubiquitylation inactive mutants failed to produce upshift bands that correspond to ubiquitylation of RNA, indicating that Ub is attached to 3’ hydroxyl group.

**Expanded View Figure 6.**
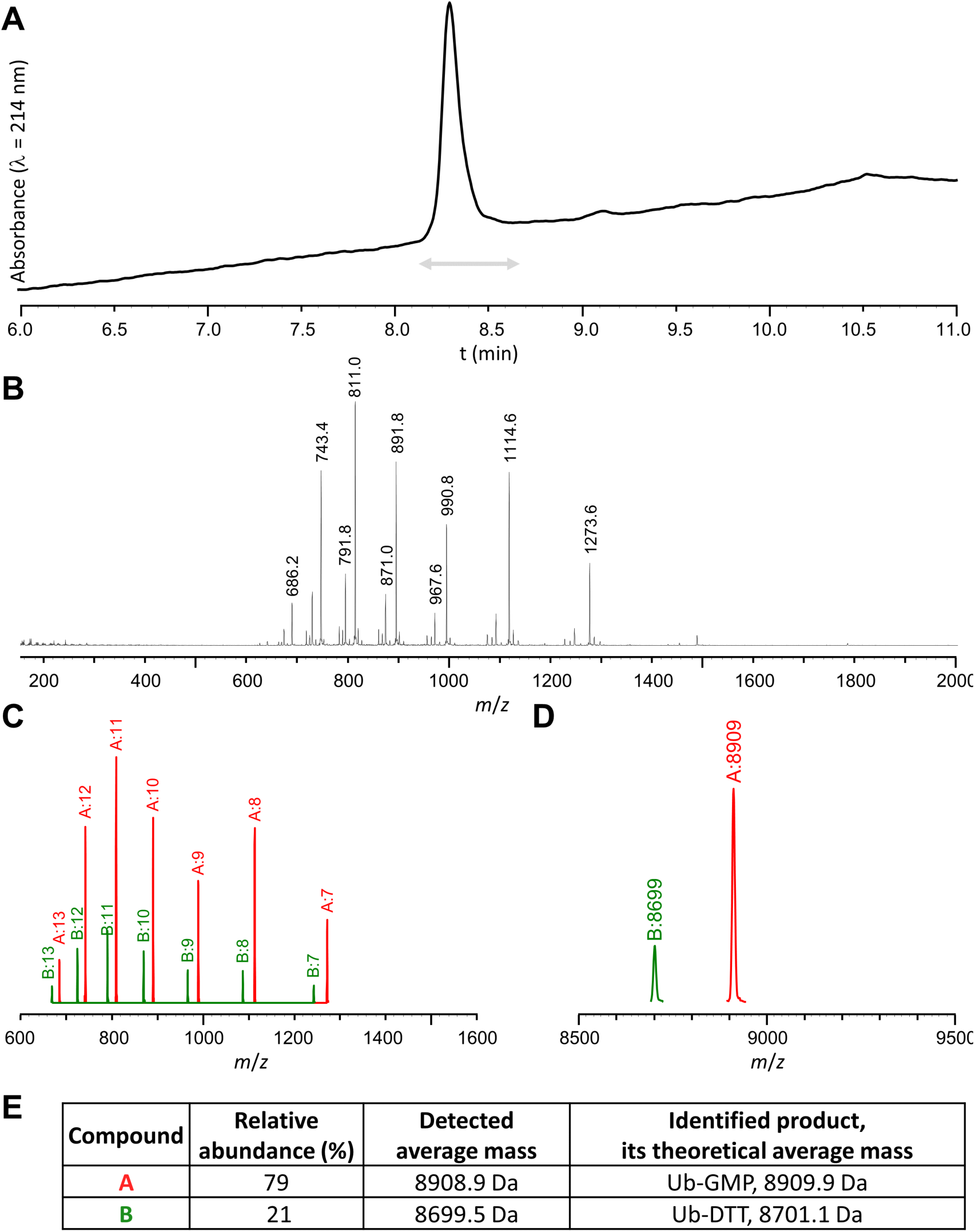
HPLC-MS analysis of the ubiquitylation mixture performed using DTX3L-RD and GMP. (A) HPLC chromatogram; (B) Experimental mass spectrum corresponding to the time window indicated as a grey arrow (sum of spectra); (C) Deconvoluted ions set, including charge state; (D) Deconvoluted spectrum; (E) Identified compounds.

**Expanded View Figure 7.**
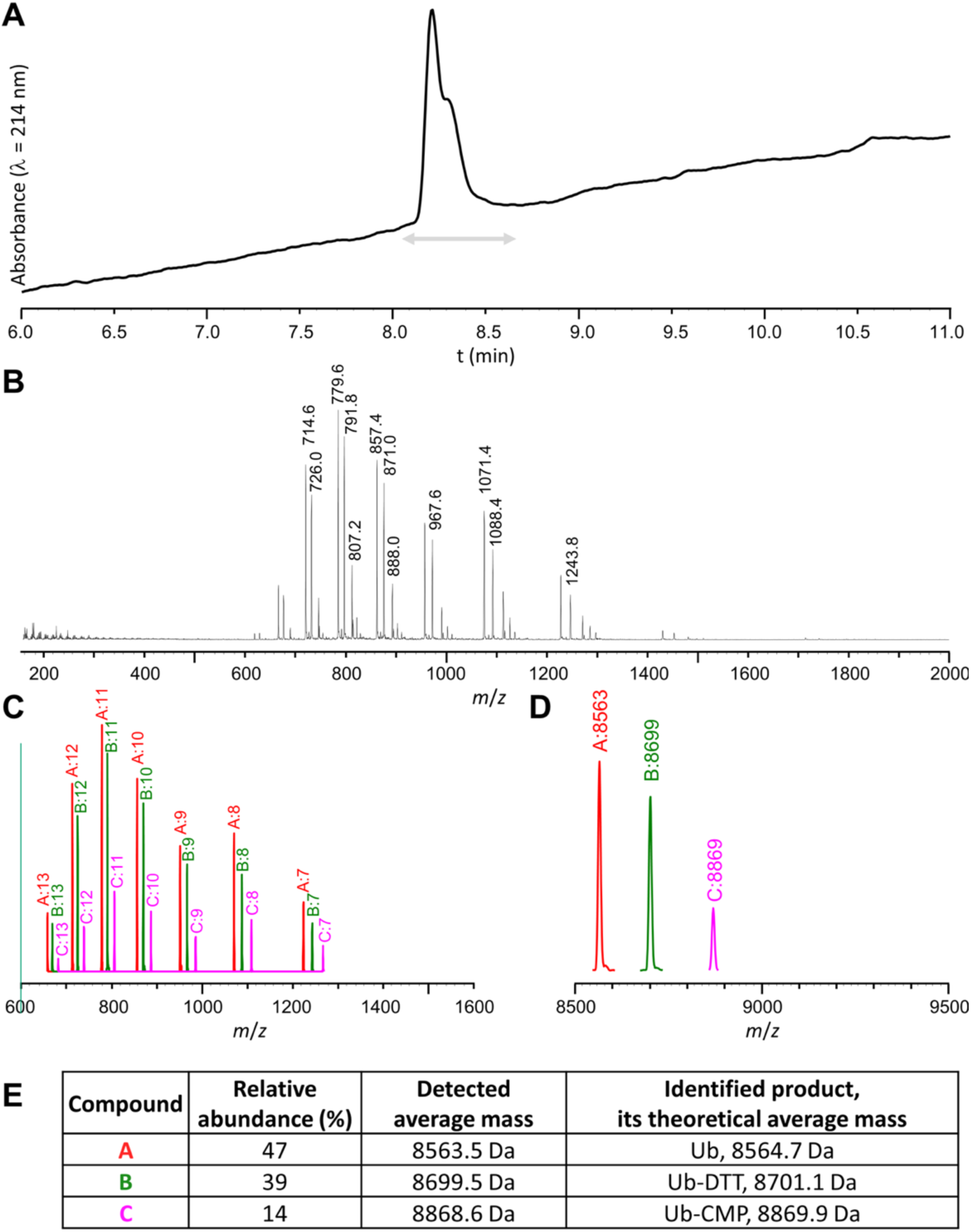
HPLC-MS analysis of the ubiquitylation mixture performed using DTX3L-RD and CMP. (A) HPLC chromatogram; (B) Experimental mass spectrum corresponding to the time window indicated as a grey arrow (sum of spectra); (C) Deconvoluted ions set, including charge state; (D) Deconvoluted spectrum; (E) Identified compounds.

**Expanded View Figure 8.**
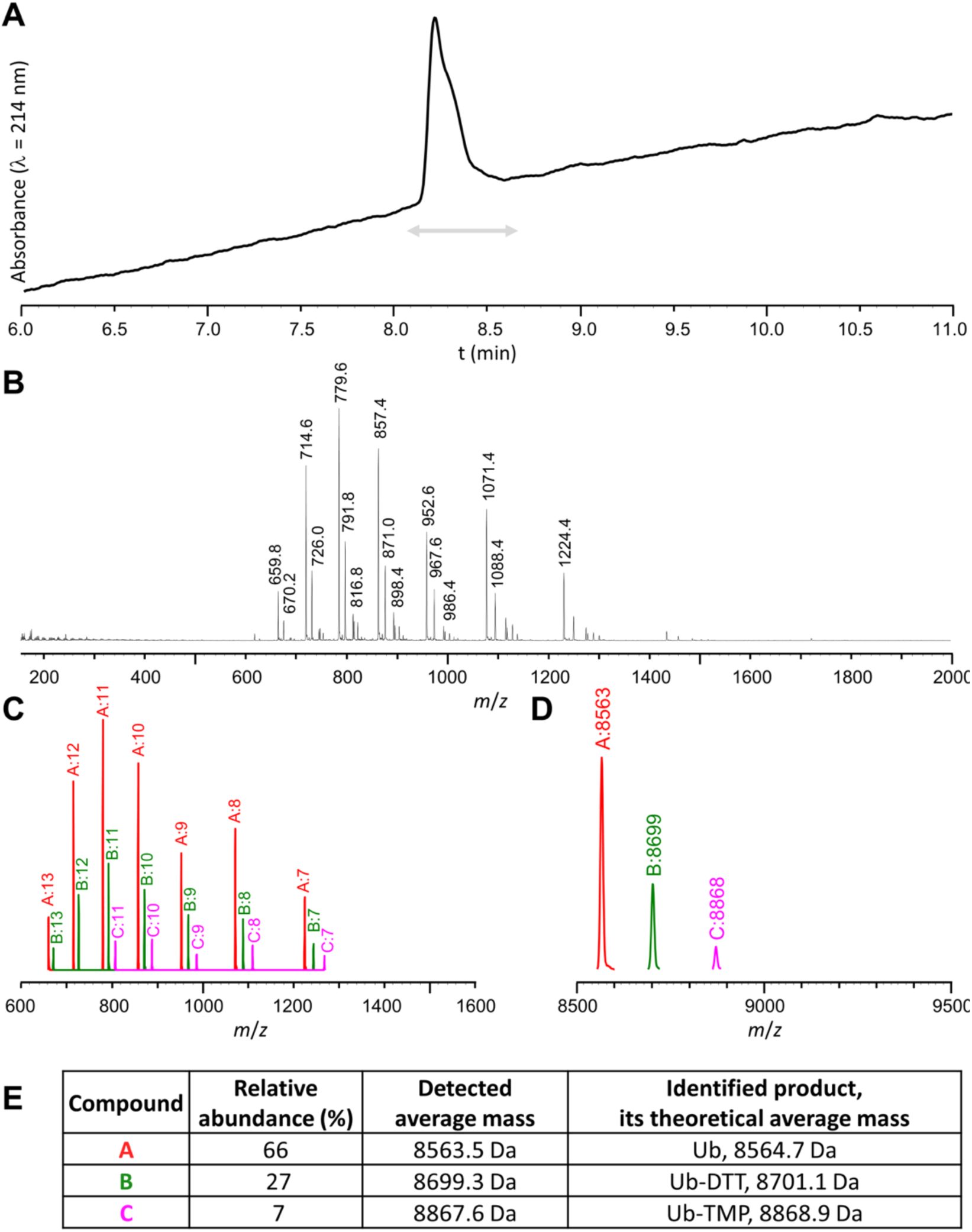
HPLC-MS analysis of the ubiquitylation mixture performed using DTX3L-RD and TMP. (A) HPLC chromatogram; (B) Experimental mass spectrum corresponding to the time window indicated as a grey arrow (sum of spectra); (C) Deconvoluted ions set, including charge state; (D) Deconvoluted spectrum; (E) Identified compounds.

**Expanded View Figure 9.**
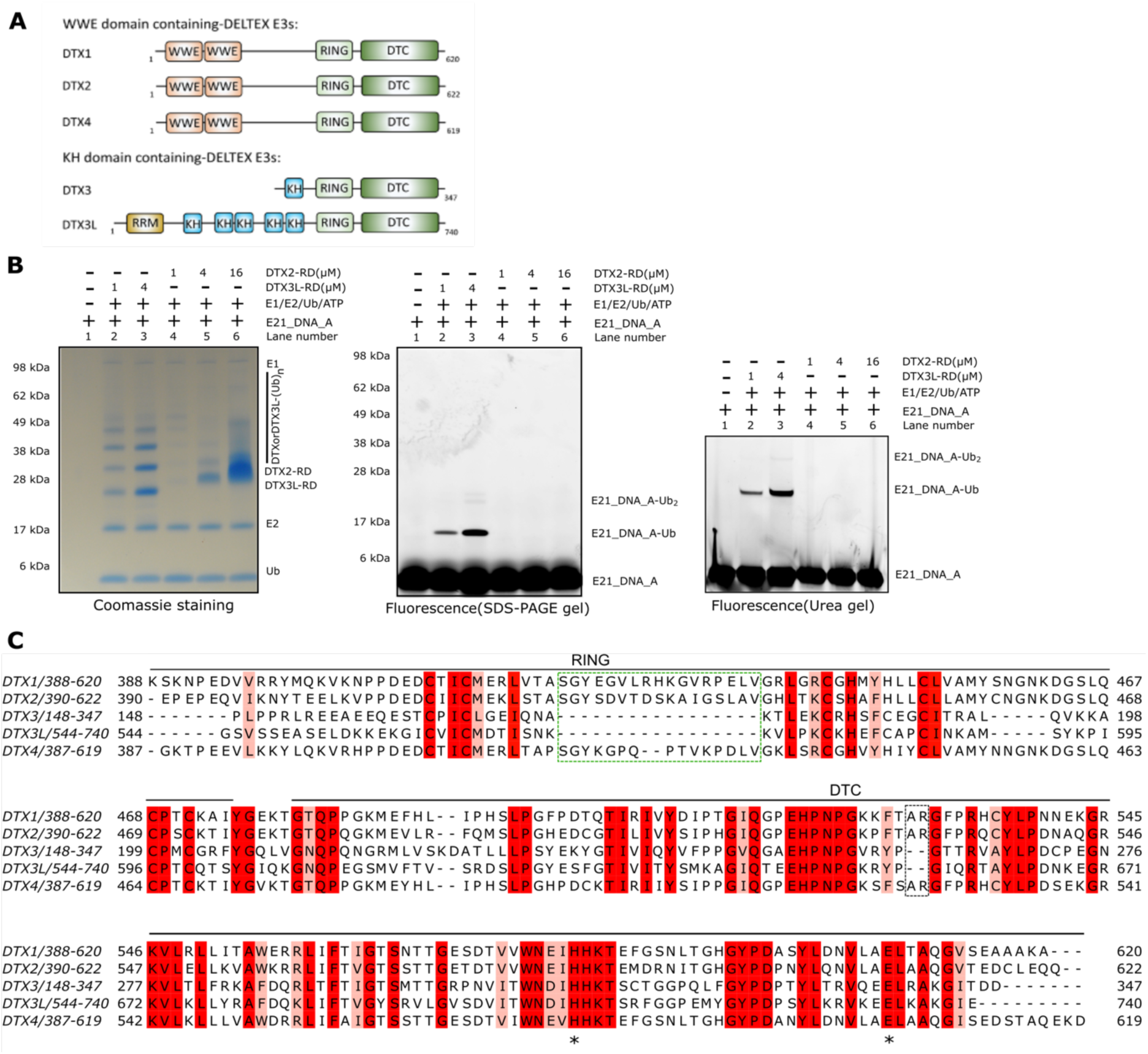
Differences between DTX2 and DTX3L. (A) Domain organisation of DELTEX family E3s. DTX1, DTX2 and DTX4 are classified into WWE domain-containing DELTEXes; DTX3 and DTX3L are classified into KH domain-containing DELTEXes. (B) DTX2-RD is not able to ubiquitylate nucleic acids. E21_DNA_A was incubated with were incubated with E1, E2, ATP, Ub and increasing amount of either DTX3L-RD or DTX2-RD, then the reactions were analysed on an SDS-PAGE gel and Urea gel and visualized using the Molecular Imager PharosFX system (BioRad). (C) Sequence alignment of the RING-DTC domains DELTEX E3s. Sequence conservation is colored in shades of red (red=identical, pink=conserved). The conserved catalytic residues in the DTC domain (DTX3L H707, E733) are indicated by asterisks. Differences in the RING domain and AR loop in DTC domain are indicated by green or black dashed box. Sequence alignment was generated using Clustal Omega (Sievers & Higgins, 2018) and Jalview (Waterhouse *et al*, 2009).

